# The Epigenetic Pacemaker is a more sensitive tool than penalized regression for identifying moderators of epigenetic aging

**DOI:** 10.1101/2021.10.05.463222

**Authors:** Colin Farrell, Kalsuda Lapborisuth, Chanyue Hu, Kyle Pu, Sagi Snir, Matteo Pellegrini

## Abstract

Epigenetic clocks, DNA methylation based chronological age prediction models, are commonly employed to study age related biology. The error between the predicted and observed age is often interpreted as a form of biological age acceleration and many studies have measured the impact of environmental and other factors on epigenetic age. Epigenetic clocks are fit using approaches that minimize the error between the predicted and observed chronological age and as a result they reduce the impact of factors that may moderate the relationship between actual and epigenetic age. Here we compare the standard methods used to construct epigenetic clocks to an evolutionary framework of epigenetic aging, the epigenetic pacemaker (EPM) that directly models DNA methylation as a function of a time dependent epigenetic state. We show that the EPM is more sensitive than epigenetic clocks for the detection of factors that moderate the relationship between actual age and epigenetic state (ie epigenetic age). Specifically, we show that the EPM is more sensitive at detecting sex and cell type effects in a large aggregate data set and in an example case study is more sensitive sensitive at detecting age related methylation changes associated with polybrominated biphenyl exposure. Thus we find that the pacemaker provides a more robust framework for the study of factors that impact epigenetic age acceleration than traditional clocks based on linear regression models.

## 1 Introduction

Epigenetic clocks, accurate age prediction models made using DNA methylation, are promising tools for the study of aging and age related biology. Beyond predicting the age of an individual to within a couple of years, multiple studies have shown that the difference between the observed and expected epigenetic age can be interpreted as a measure of biological age acceleration [1]. Age acceleration observed using the first generation of epigenetic clocks [2, 3] has been associated with a variety of health outcomes including mortality risk[4, 5], cancer risk [6], cardiovascular disease[7] and other negative health outcomes[8–10]. However, as epigenetic clocks become more accurate, epigenetic age acceleration is no longer associated with mortality [11].

Epigenetic clocks are generally trained using a regularized regression model. Given an elastic net model of the form *y* = *βX* the goal of penalized regression is to maximize the likelihood by reducing the prediction error of the model, *L*(*λ*_1_, *λ*_2_, *β*) = |*y* − *Xβ*|^2^ + *λ*_2_|*β*|^2^ + |*λ*_1_*β*|. In the case of epigenetic clocks, the likelihood is maximized by minimizing the difference between the observed and predicted age subject to the elastic net penalty,*λ*_1_ and *λ*_2_.. Methylation sites that increase modeled error but contain biologically meaningful information may be discarded during model fitting. This problem is magnified in the case of epigenetic clocks where the relationship between methylation and time is nonlinear[12].

An alternative and complementary approach to studying epigenetic aging is to model how methylation changes for a predetermined collection of sites with respect to time. To this end, we have developed the epigenetic pacemaker (EPM) [13, 14] to model methylation changes with age. Given *j* individuals and *i* methylation sites, under the EPM an individual methylation site can be modeled as 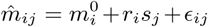 where 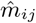 is the observed methylation value, 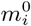 is the initial methylation value, *r*_*i*_ is the rate of change, *s*_*j*_ is the epigenetic state, and *ϵ*_*ij*_ is a normally distributed error term. The *r*_*i*_ and 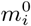 are characteristic of the sites across all individuals and the epigenetic state of an individual *s*_*j*_ is set using information from all modeled sites. Given an input matrix 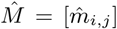 the EPM utilizes a fast conditional expectation maximization algorithm to find the optimal values of 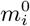, *r*_*i*_, and *s*_*j*_ to minimize the error between the observed and predicted methylation values across a set of sites. This is accomplished by first fitting a linear model per site using age as the initial *s*_*j*_. The *s*_*j*_ of the modeled samples is then updated to minimize the error between the observed and predicted methylation values. This process is performed iteratively until the reduction in error is below a specified threshold or the maximum number of iterations is reached. Under the EPM, the epigenetic state has a linear relationship with the modeled methylation data, but not necessarily with chronological age. This allows for nonlinear relationships between time and methylation to be modeled without prior knowledge of the underlying form. Every modeled methylation site has a characteristic 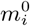 and *r*_*i*_ that describes the site in relation to other modeled sites and the output epigenetic states. In the current work, we ask whether the EPM formalism can be utilized for the identification of moderators that impact the association between age and epigenetic state (i.e factors that accelerate or decelerate the changes in epigenetic states with time). To this end we extend the EPM model to simulate methylation matrices associated with age and age accelerating phenotypes. We then evaluate the ability of regularized regression and EPM models to detect age acceleration traits that have linear and nonlinear associations with age. Utilizing a large aggregate data set we validate the simulation results and in one illustrative example further assess the ability of both approaches to detect age related methylation changes associated with PBB exposure.

## 2 Results

### 2.1 Simulation of Trait Associated Methylation Matrices

Under the EPM the epigenetic state for individual *j, S*_*j*_, can be interpreted as a form of biological age that represents a weighted sum of aging associated phenotypes 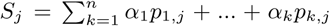. Under this model *α*_*k*_ is the weight for phenotype *k* and *p*_*k,j*_ is the value of phenotype *k*. Phenotypes may contribute to increased or decreased aging respectively and when considered as a whole contribute to the overall aging rate observed for an individual.

As shown in our previous work[12], the relationship between *p*_*k,j*_ and time is not necessarily linear. When simulating age associated phenotypes, each phenotype can be represented as 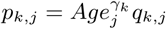, where *γ*_*k*_ is a phenotype specific parameter shared among all individuals and *q*_*j,k*_ represents the magnitude of exposure for a simulated trait and is personal to an individual. The observed phenotype is modeled as an interaction between age and an exposure of varying magnitude among individuals. This formulation is flexible as non-age dependent traits can be easily simulated by setting 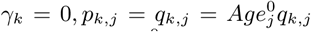. Individual sites can be described as a linear model where 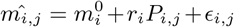. *P*_*i,j*_ is a weighted sum of phenotypes influencing the methylation status of an individual site, 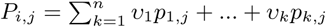.

To assess the sensitivity of the EPM and penalized regression approaches at detecting moderator of epigenetic aging we simulated a methylation matrix containing linear and nonlinear age associated traits of form 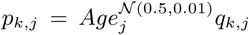 and 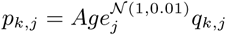. The trait *γ* parameter was generated by sampling from a normal distribution 𝒩(0.5, 0.01) to generate traits with varying relationships with time (Figure 1). Samples were simulated by assigning an age from a uniform distribution, 𝒰(0, 100) and setting sample health by sampling from a normal distribution. Sample health is a sample specific metric that influences the magnitude and direction of the simulated age accelerating trait. Simulated traits included a binary phenotype (*P* = 0.5), continuous phenotypes influenced by only age, or by age and sample health (Table 1). The effect, *q*, of a binary trait was varied from 0.995 to 1.0 over 5 equally spaced intervals. Given a binary trait form of 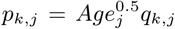 a 0.001 decrease in *q* corresponds to a 1 percent decrease in epigenetic state by age 100 relative to samples not assigned the binary trait. Within each interval the standard deviation of the sample health sampling distribution was varied from 0.0 to 0.01 over 5 equally spaced intervals. The simulation was repeated 50 times for each binary, continuous trait combination with 500 simulated samples within each simulation. Additionally, at a binary *q* of 0.995 the range of continuous traits was expanded over a broader range to assess the model sensitivity for detecting the continuous trait. Five methylation sites for all continuous traits were then simulated and 50 methylation sites for the binary trait. An additional 50 sites were simulated that were equally influenced by a mixture of four continuous traits and the simulated binary trait. The resulting simulation matrix contains 450 methylation sites.

**Figure1:**
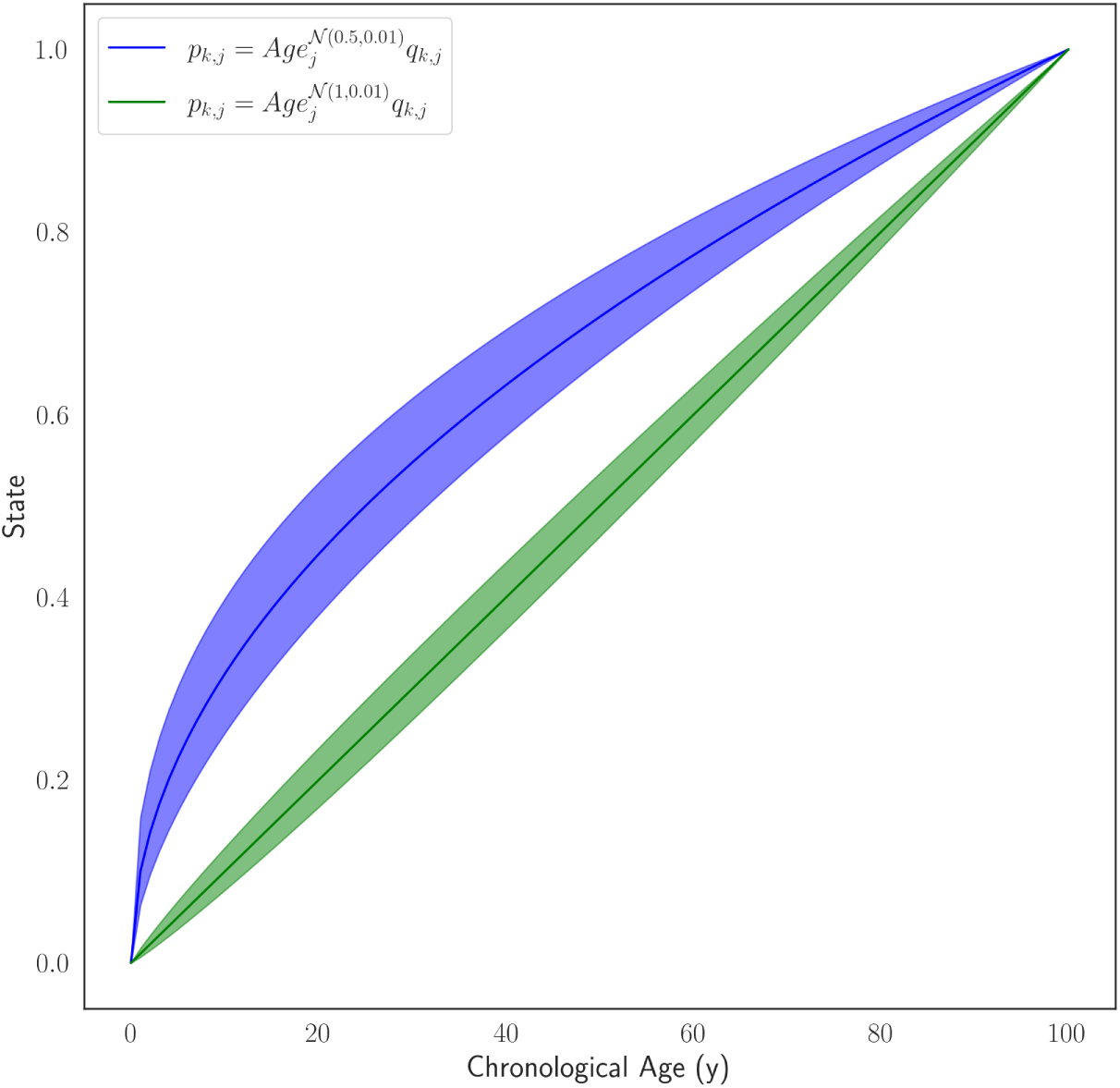
Simulated trait forms where the shaded area represent one standard deviation away from the mean *γ*, given 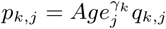.

**Table 1:** Simulated Trait Conditions

| Trait Form | Beta | Gamma | Gamma Std. Dev. | Sample Effect | Age Only | Generated Phenotypes |
| --- | --- | --- | --- | --- | --- | --- |
| Continuous | 0.1 | $\mathcal{N}(0.5, 0.01)$ | 0.05 | Yes | No | 10 |
| Continuous | 0.1 | $\mathcal{N}(1.0, 0.01)$ | 0.05 | Yes | No | 10 |
| Continuous | 0.1 | $\mathcal{N}(0.5, 0.01)$ | 0.05 | No | Yes | 20 |
| Continuous | 0.1 | $\mathcal{N}(1.0, 0.01)$ | 0.05 | No | Yes | 20 |
| Binary<br>($Pr = 0.5$) | 0.1 | 0.5 | 0 | Yes | No | 1 |

Given a simulation data set, the samples were split randomly in half for model training and testing. EPM and penalized regression models were fit for each simulation training set and epigenetic state and age predictions were made for the testing set. e then fit a regression model where the epigenetic age or state is dependent on the age, square-root of the age, the health status, and binary trait status of the sample 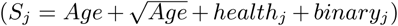. The square-root of the age is included in the regression model to account for the nonlinear relationship between the simulated age and methylation data.

As the exposure size of the binary trait is decreased from 1.00 to 0.995 the ability to detect the influence of the trait on the epigenetic state and age is improved (Figure 2A and B). At an effect size of 0.995 the estimated effect of the binary trait on the epigenetic state is significant (*µ* = 0.035, *σ* = 0.089) while the effect on the epigenetic age it is not (*µ* = 0.269, *σ* = 0.282). At an exposure size of 1.0, where the simulated binary trait has no effect, the distribution of p values forEPM and linear models is randomly distributed. The ability to observe the health effect of the simulated continuous traits improves in both the linear and EPM models as the standard deviation of the sample health sampling distribution is increased (Figure 2 C and D). At an exposure size of 0.002 and 0.0025 the average EPM model is significant (*µ* = 0.0194, *σ* = 0.0436) while the average linear model is not (*µ* = 0.0607, *σ* = 0.128). At a continuous trait standard deviation above 0.005 both models produce significant results.

**Figure2:**
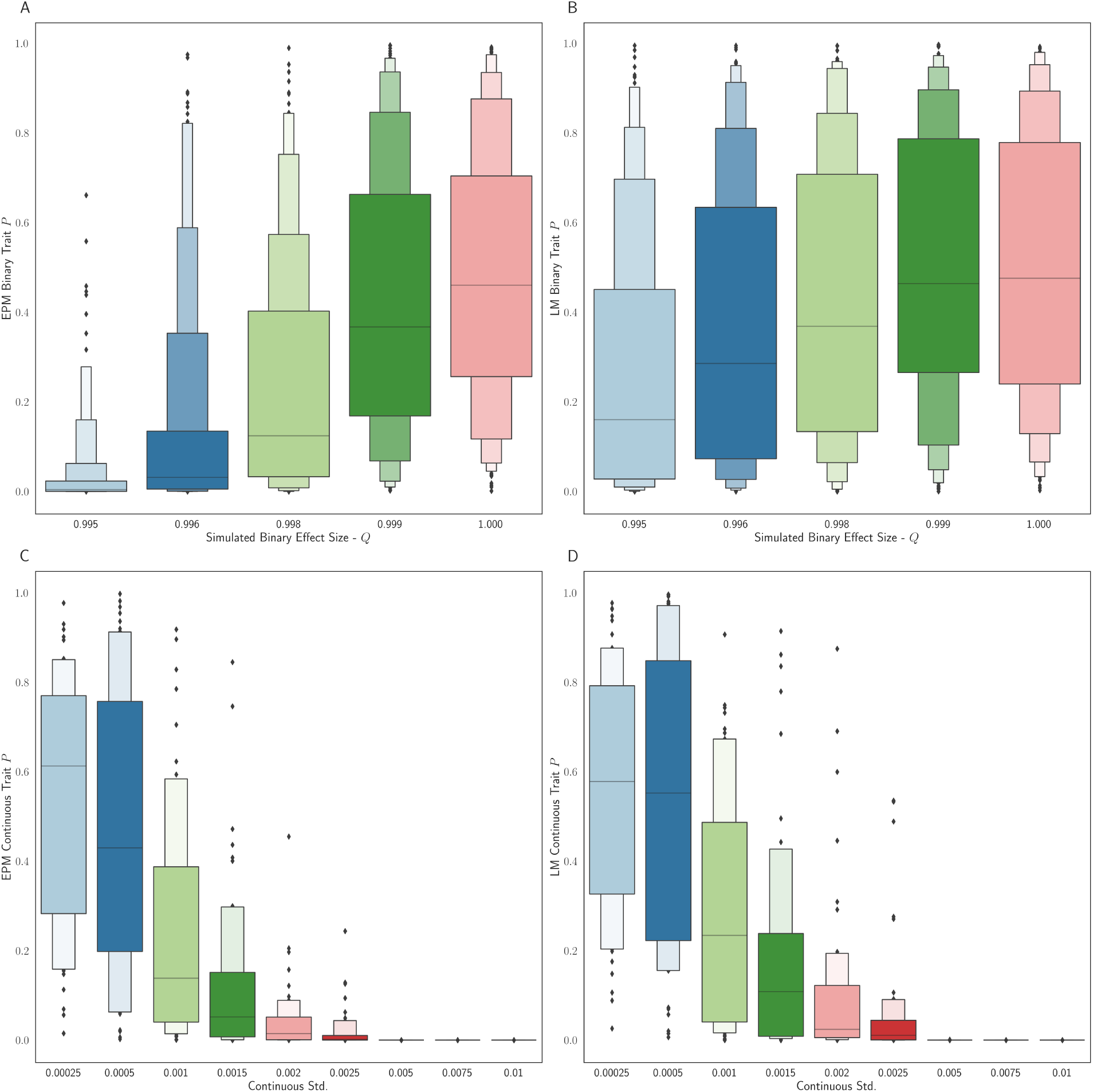
The distribution binary coefficient p-values for **A** EPM and **B** penalized regression models. The distribution of p-values given a simulation health standard deviation for **C** EPM and **D** penalized regression models.

### 2.2 Universal Blood EPM and Penalized Regression Models

We validated the simulation results using a large aggregate data set composed of Illumina 450k array data[15–27] deposited in the Gene Expression Omnibus[28] (GEO). All methylation array data sets were processed using a unified pipeline from raw array intensity data (IDAT) files using minfi (Aryee et al., 2014). Sex and blood cell type abundance predictions were made for each processed as previously described[29, 30]. The aggregate data set contains 6,251 whole blood tissue samples representing 16 GEO series.

We trained EPM and penalized regression models using data assembled from four GEO series[31–34] (*n* = 1605) with samples spanning a wide age range (0.01 - 94.0 years). The training set was split by predicted sex, within each sex we used stratified sampling by age to select 95% of the samples for model training. The selected samples from each sex were combined (*n* = 1524) and the remaining samples (*n* = 81) left out for model evaluation. Methylation values for all samples were quantile normalized by probe type[2] using the median site methylation values across all training samples for each methylation site. Principal component analysis (PCA) was performed on the cell type abundance estimates using the training data. The trained PCA model was used to predict the cell type PCs for the testing and validation data sets.

We fit a penalized regression model to the training matrix as follows. The normalized training methylation matrix was first filtered to remove sites with a variance below 0.001, resulting in a training matrix with 183,114 sites. A cross validated (*cv* = 5) elastic net model was trained against training sample ages using the filtered methylation matrix. The trained model performed well on the training (*R*^2^ = 0.981) and testing (*R*^2^ = 0.940) data sets (S.Figure 2).

In contrast to penalized regression based approaches, site selection for the EPM model is performed outside of model fitting. Methylation sites were selected for model training if the absolute Pearson correlation coefficient between methylation values and age was greater than 0.4 (*n* = 16, 880). A per site regression model was fit using the observed methylation value as the dependent variable and age as the explanatory variable. Sites with a mean absolute error (MAE) less than 0.025 between the predicted and observed methylation values were retained for further analysis (*n* = 7, 013). An EPM model was fit using these sites (Figure 3A). We then sought to identify subsets of sites that had functionally similar forms between age and methylation. This was done to filter sites that were associated with age by chance and to select clusters of sites with low prediction error. Subsets of sites with similar functional form were identified by clustering sites using affinity propagation [35]) by the euclidean distance between the single site regression model residuals. Cross validated EPM and penalized regression models were trained for all clusters with greater than ten sites (*n* = 55). The cluster EPM models show varying associations between the epigenetic state and age relative to the EPM model fit with all sites initially selected by absolute PCC(Figure 3B). Clusters with an observed EPM and penalized regression MAE less than 6 (*n* = 5) were combined to fit final EPM and penalized regression models. This resembles the simulated methylation matrices where sites with differing functional forms are modeled collectively. The combined cluster EPM and combined cluster regression model performed well on the training and testing data sets (S.Figure 1).

**Figure3:**
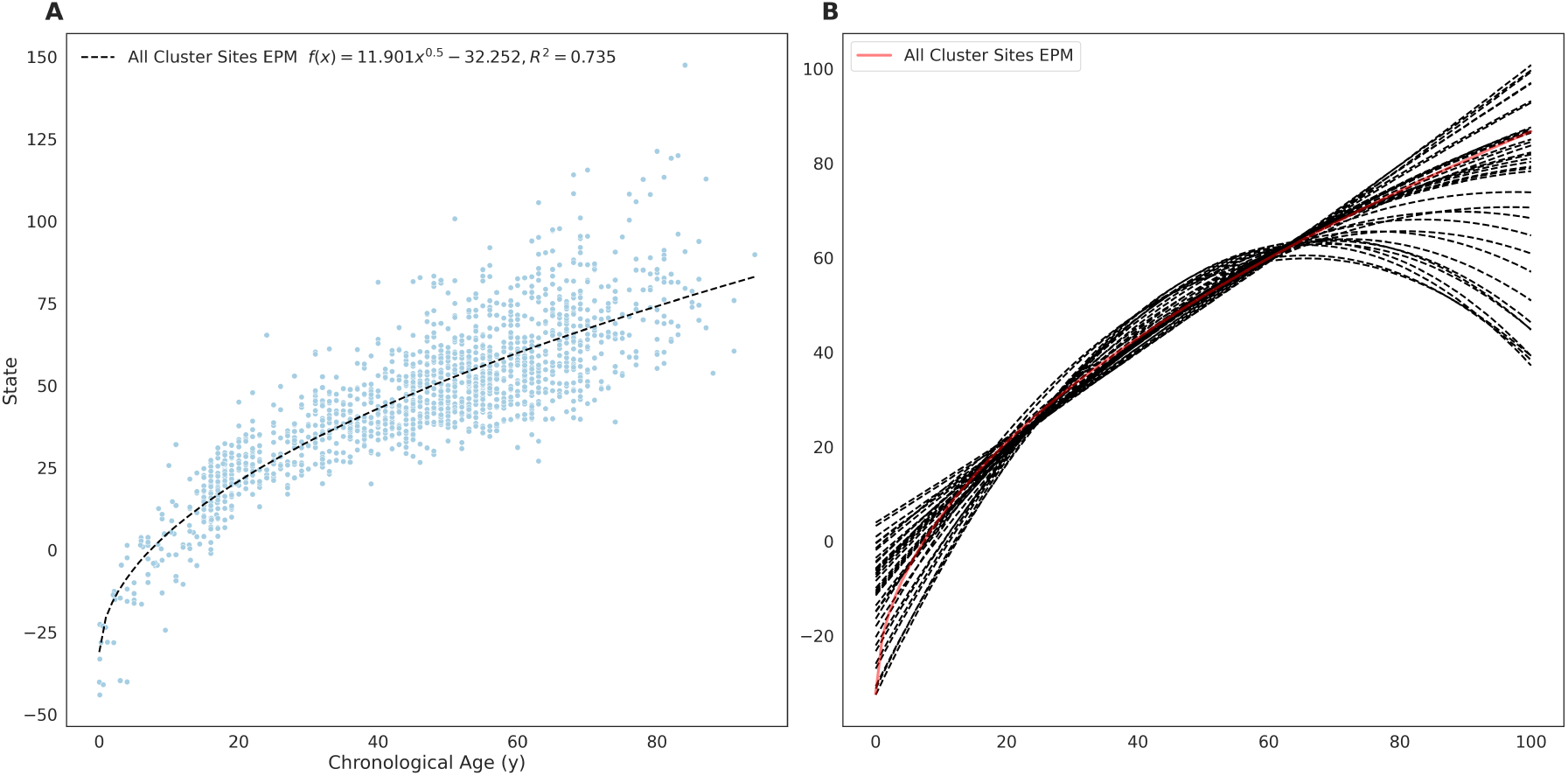
**A** EPM model fit with 3832 methylation sites with a MAE below 0.025. **B** The fit trend line for EPM clusters with more than 10 sites and an *R*^2^ ≥ 0.4.

We evaluated the combined cluster EPM, combined cluster penalized regression, and the full penalized regression models against a validation data set consisting of 14 GEO series experiments representing 4,600 whole blood tissue samples. Each model accurately predicted the epigenetic state or epigenetic age of the validation samples (Figure 4). We then fit an ordinary least squares regression model for every validation experiment individually to predict the observed epigenetic age or state using the sample age, the square root of age, cell type PCs, and predicted sex 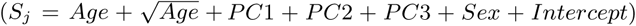. If the proportion of female samples to the total number of samples was greater than 0.7 the sex term was dropped from the regression model. Significant cell type PC2 coefficients were observed for all EPM models and the majority of the cluster and full penalized regression models (Figure 5A). Significant cell type PC1 and PC3 coefficients were observed for the majority of the EPM models but not for the cluster or full penalized regression models. Significant sex effects (*p <* 0.0038) were observed for 9, 4, 0 out of 15 models for the EPM, cluster penalized regression, and full penalized regression respectively (Figure 5B).

**Figure4:**
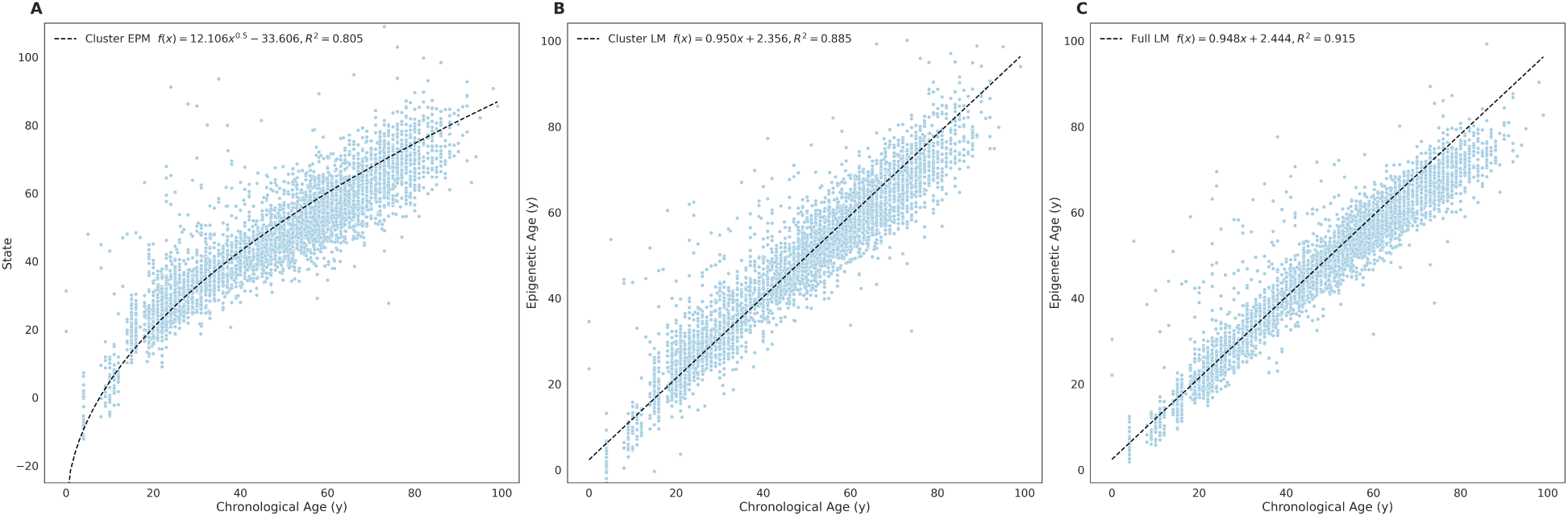
Whole blood tissue validation **A** EPM, **B** cluster penalized regression and **C** full penalized regression models.

**Figure5:**
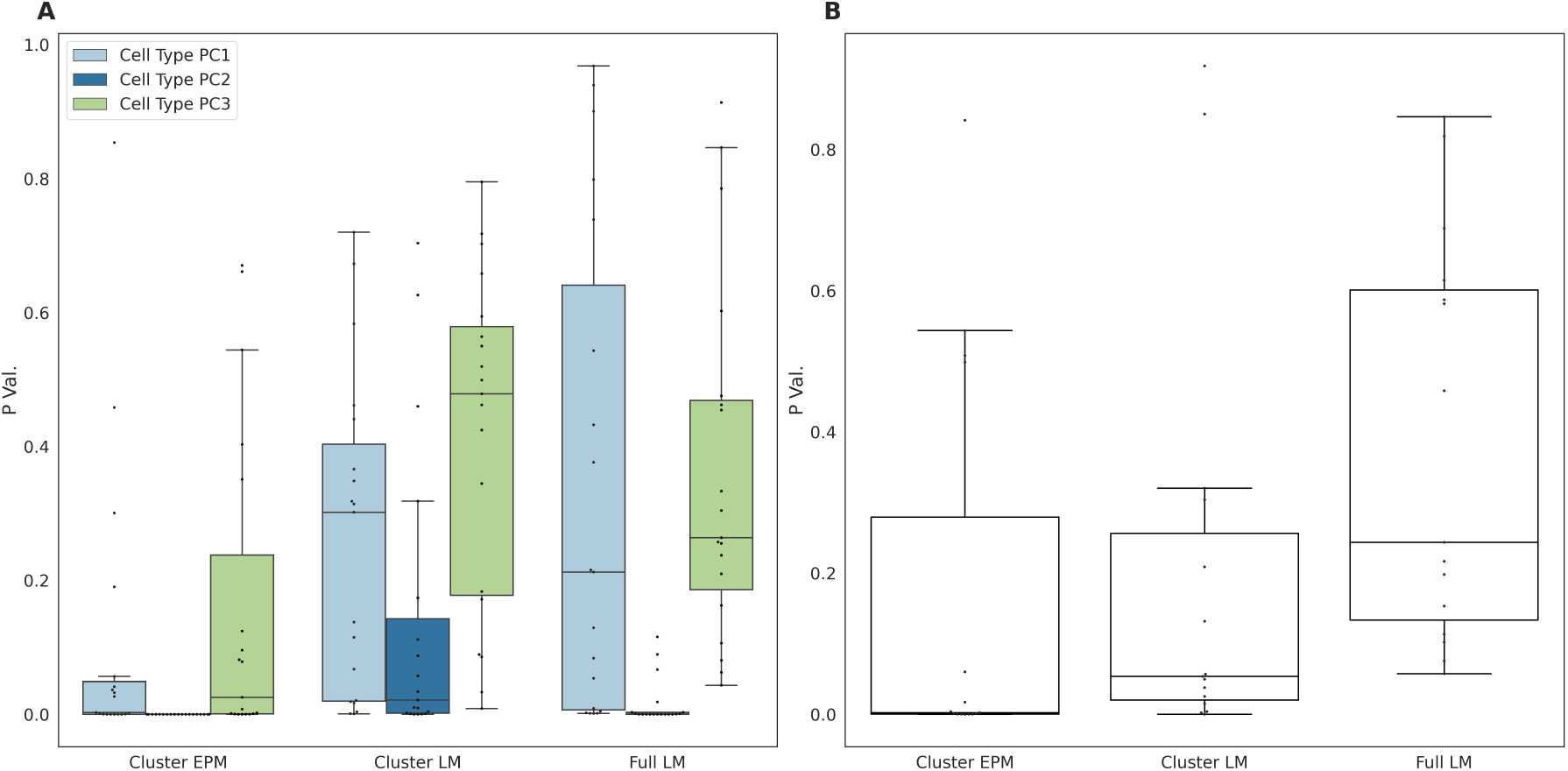
**A** Cell type principal component and **B** predicted sex regression coefficient p-values.

### 2.3 Polybrominated Biphenyls Exposure

Polybrominated biphenyls (PBB) were widely used throughout the United States in the 1960’s and 1970’s for a variety of industrial applications. Widespread PBB exposure occurred in the state of Michigan from the summer of 1973 to later spring of 1974 when an industrial PBB mixture was incorrectly substituted for a nutritional supplement used in livestock feed[36]. PBB is biologically stable and has a slow biological half life; individuals exposed during the initial 1973 - 1974 incident still have detectable PBB in their blood[37]. PBB is an endocrine-disrupting compound and exposure has been linked to numerous adverse health outcomes in Michigan residents such as thyroid dysfunction[38, 39] and various cancers[40, 41]. A study by Curtis et al. showed total PBB exposure is associated with altered DNA methylation at CpG sites enriched for an association with endocrine-related autoimmune disease [42]. Utilizing the publicly available Illumina Methylation EPIC array [43] profiles (*n* = 679), that covered a wide age range (23 - 88 years), we sought to compare the ability of penalized regression and the EPM to detect epigenetic age acceleration associated with PBB exposure.

Briefly, 50% of samples (*n* = 339) were selected for model training using stratified cross validation by age. A cross validated elastic net model was trained using all methylation sites with a site variance above 0.001, (*n* = 529, 703). The trained model performed well on the training and testing data sets (*R*^2^ = 1.00, *R*^2^ = 0.740, *S.Figure*2*A* − *B*). EPM sites were selected and models fit as described with the universal blood EPM. Four EPM clusters (*MAE <* 6) were merged for a combined EPM model built using 413 CpG sites. The combined EPM model performed well in training and testing data sets (*R*^2^ = 0.794, *R*^2^ = 0.812, *S.Figure*2*C* − *D*). Epigenetic age and epigenetic state predictions were then made for the testing samples using the penalized regression and EPM models.

We then fit an OLS regression model to predict the epigenetic age or state dependent on PBB-153 exposure, h age, the square root of age, cell type PCs, and predicted sex 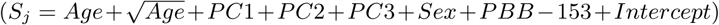. PBB-153 exposure was highly significant in the EPM regression model (*p* = 5.9*e* − 10) but not the penalized regression model (*p* = 0.141).

## 3 Discussion

A long standing question in the field of epigenetics was whether biomarkers could be trained to predict various traits using methylation measurements. The most successful biomarkers to date have been epigenetic clocks that can accurately predict the age of an individual based on their methylation pattern. These have been shown to be useful for human studies of aging, as well as for animal studies, including mice[44] and dogs[45]. DNA methylation biomarkers are typically constructed using penalized regression approaches. Given a large enough matrix, penalized regression will select sites that minimize the prediction error given a modeled trait. Epigenetic clocks are examples of such models. Beyond predicting actual ages, these models have also been used to measure the influence of external factors on the rates of aging, and multiple studies have shown that the resulting age accelerations (i.e the differences between actual and predicted ages) are significantly associated with multiple factors such as cardiovascular disease[7] and mortality risk[4, 5].

While epigenetic clocks have proven to be useful they have significant limitations. Because they are based on linear models, it may be difficult to model aging when the underlying methylation changes are non-linear in time. Moreover, epigenetic clocks are prone to over fitting, and while cross validation schemes are often used to construct robust clocks, they often do not yield accurate estimates for other data sets. Finally, epigenetic clocks are not very interpretable, and highly degenerate, so that it is difficult to extract biological insights from the weights of the models.

To overcome some of these limitations, we have previously developed the epigenetic pacemaker formalism. In this approach, rather than building a model for the age, we construct a model for the observed methylation data that depends on age. The advantage of this approach is that this formalism allows us to identify non-linear associations between methylation and age across a lifespan. Moreover, these models tend to be robust to training as they are fit to large methylation matrices rather than age vectors. Finally, the model describes the change in methylation at each site with respect to a time dependent epigenetic state, and therefore all parameters of the model are directly interpretable as either initial values of methylation or rates of change of methylation.

Depending on the context, epigenetic clocks are both more and less effective than the EPM. The penalized regression models provide more accurate age predictions (*R*^2^ = 0.875, 0.911) than the EPM model (*R*^2^ = 0.821), and the model output can be directly compared to the age of a sample. However, because these models are optimized to reduce the error between actual and predicted age, they tend to minimize the effect of extraneous factors on the predicted age. As such, epigenetic clocks are not optimal for identifying external factors that moderate the relations between actual and predicted age. By contrast, the EPM models are not optimized to minimize the difference between predicted and actual age, but rather try to minimize the difference between observed and modeled methylation values. As such, they retain many of the effects that other factors may have on the association between methylation and epigenetic states.

In this study we find that while the penalized regression models were more accurate for predicting age, the epigenetic state generated by the EPM is significantly impacted by cell type and sex effects in both simulations and real data. We also found that The EPM model generated for individuals exposed to PBB was sensitive to e PBB exposure, which has been linked to negative health outcomes, while the penalized regression epigenetic aging model was not. Additionally, the sensitivity of the EPM to moderators of epigenetic aging has been supported by two two recent studies investigating epigenetic aging in marmots[46] and zebras[47]. In the first of these studies, EPM models showed an association between hibernation and slowed epigenetic aging in marmots and in the second an increased epigenetic age associated with zebra inbreeding; no such associations were observed with penalized regression epigenetic age models.

Most studies of human epigenetic aging are not motivated by the development of accurate age predictors, since ages are nearly always known in studies, but rather by the discovery of biological aging moderators. The EPM is a more sensitive approach than epigenetic clocks for the detection of factors other than age that influence the epigenome and therefore potentially more useful for discovering moderators of biological aging.

## 4 Methods

### 4.1 Simulation

We implemented the simulation framework as a python package with numpy(≥v1.16.3)[48] and scikit-learn(v0.24)[49] as dependencies. A simulation run generates a trait-associated methylation matrix and samples are tied to the simulated traits. The simulation procedure is implemented as follows:

- Traits are intialized that contain the information about the trait relationship with age and a simulated sample phenotype. Given the structure 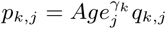, and *k* samples and *j* traits *γ* is characteristic of the trait. When a sample is passed to a trait, a value of *q* is generated for the sample by sampling from a normal distribution with a variance characteristic of the simulation trait. Additionally, each trait can be optionally influenced by a characteristic measure of sample health, *h*_*j*_. Given, a normally distributed trait 𝒩(*µ, σ*^2^) and a health effect *h*_*j*_, the sampled distribution for individual *j* is 𝒩(*µ* + *h*_*j*_, *σ*^2^). Continuous and binary traits can be simulated. If a binary trait is simulated, a *q* other than 1 is assigned at a specified probability.
- Samples are simulated by setting the age by sampling from a uniform distribution over a specified range and by setting a sample health metric *h* by sampling from a normal distribution centered on zero with a specified variance. Traits passed to a sample simulation object are then set according to the age and health of the sample. Simulated samples retain all the set phenotype information for downstream reference.
- Methylation sites are simulated by randomly setting the initial methylation value, maximum observable methylation value, the rate of change at the site, and the error observed at each site. Sites are then assigned traits that influence the methylation values at each site.
- Methylation values are simulated for each site for every individual given the simulated phenotypes with a specified amount of random noise.

### 4.2 Simulation EPM and Penalized Regression Models

Simulation data was randomly split in half into training and testing sets. EPM models were fit using the simulated methylation matrix against age. Penalized regression models were fit using scikit-learn(v0.24)[49] ElasticNet (alpha=1, l1 ratio=0.75, and selection=random). All other parameters were set to their default values. Ordinary least squares regression as implemented in statsmodels (0.11.1)[50] was utilized to describe the epigenetic state or age with the following form 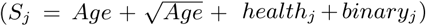. Full analysis is found in the EPMSimulation.ipynb supplementary file.

### 4.3 Methylation Array Processing

Metadata for Illumina methylation 450K Beadchip methylation array experiments deposited in the Gene Expression Omnibus (GEO) [28] with more than 50 samples were parsed using a custom python tool set. Experiments that were missing methylation beadchip array intensity data (IDAT) files, made repeated measurements of the same samples, utilized cultured cells, or assayed cancerous tissues were excluded from further processing. IDAT files were processed using minfi[30] (v1.34.0). Sample IDAT files were processed in batches according to GEO series and Beadchip identification. Methylation values within each batch were normal-exponential normalized using out-of-band probes[51]. Blood cell types counts were estimated using a regression calibration approach[29] and sex predictions were made using the median intensity measurements of the X and Y chromosomes as implemented in minfi[30]. Whole blood array samples were used for downstream analysis if the sample median methylation probe intensity was greater than 10.5 and the difference between the observed and expected median unmethylation probe intensity is less than 0.4, where the expected median unmethylated signal is described by (*y* = 0.66*x* + 3.718).

### 4.4 Blood EPM and Penalized Regression Models

Methylation sites with an absolute Pearson correlation coefficient between methylation values and age greater than 0.40 and 0.45 for the unified whole blood and PBB data sets respectively were initially selected for EPM model training. A linear model was generated using numpy polyfit [48] with age and the independent variable and methylation values as the dependent variable. Mean absolute error (MAE) was calculated as the mean absolute difference between the observed and predicted meth values according to the site linear models. A vector of residuals generated using these models were utilized for clustering by affinity propagation[35]) as implemented in scikit-learn (v0.24)[49] with a random state of 1 and a cluster preference of -2.5. Cross-validated EPM, and penalized regression models for the universal blood analysis, were trained for all clusters containing greater than ten sites. Clusters with an observed EPM and penalized regression MAE less than 6.0 were combined to fit final EPM and regression models.

Penalized regression models were fit using scikit-learn(v0.24)[49] ElasticNetCV (cv=5 alpha=1, l1 ratio=0.75, and selection=random). All other parameters were set to their default values. Principal Component Analysis as implemented in scikit-learn was utilized with default parameters to perform PCA on training sample cell type abundances. The trained PCA was utilized to calculate cell type PCs for the testing and validation samples. Ordinary least squares regression as implemented in statsmodels (0.11.1)[50] was utilized describe the epigenetic state or age with the following form 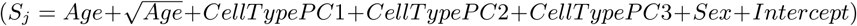. Full analysis is found in the EPMUniversalClock.ipynb supplementary file.

### 4.5 Analysis Environment

Analysis was carried out in a Jupyter[52] analysis environment. Joblib[53], SciPy[54], Matplotlib[55], Seaborn[56], Pandas[57] and TQDM[58] p ackages were utilized during analysis.

## 4.6 Supplementary Information

Analysis code and notebooks can be found at https://github.com/NuttyLogic/EPM-ModeratorsOfAging.

---

**S.Figure1:**
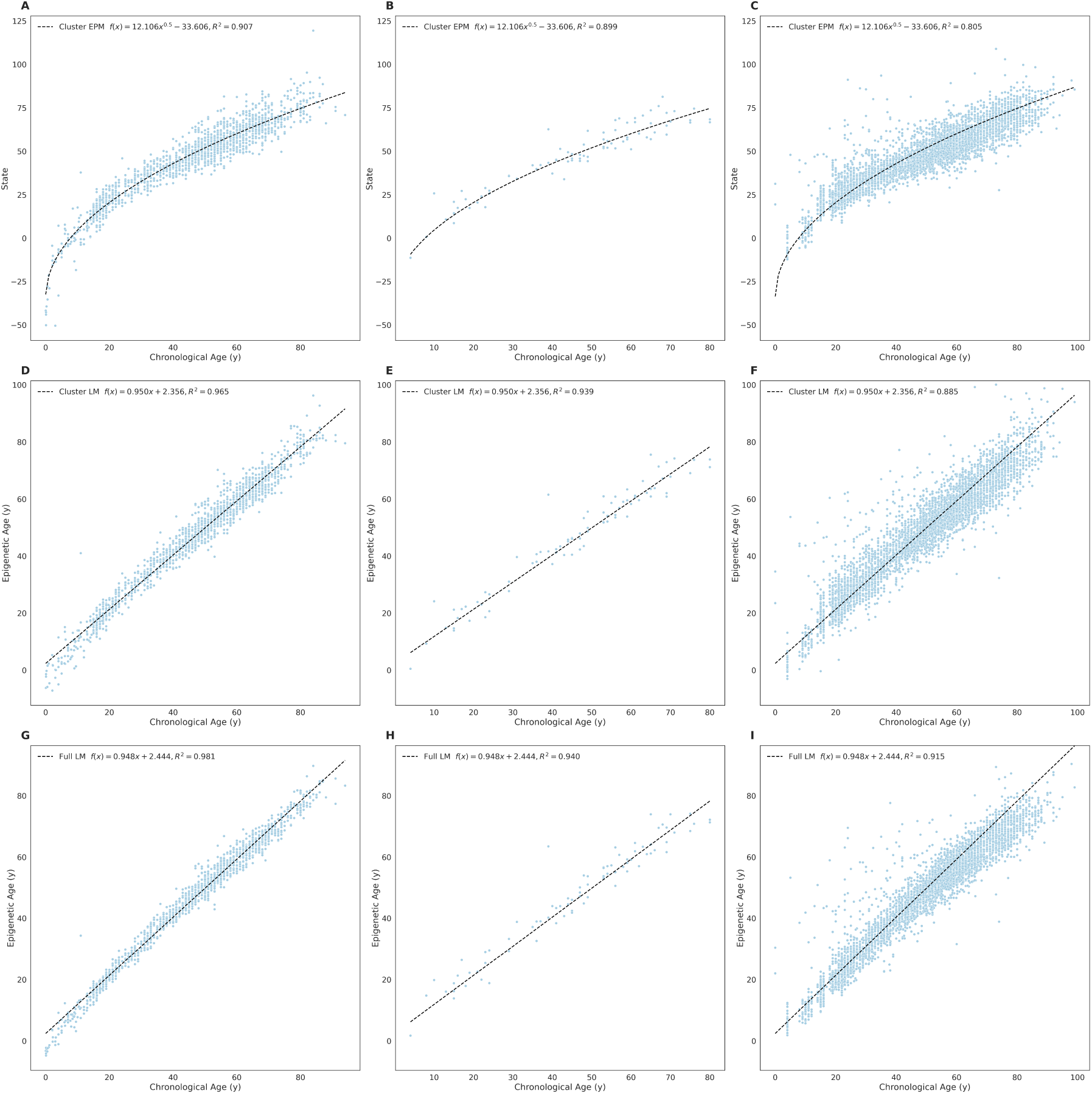
Universal blood EPM and regression models. **A - C** Train, testing, and validation EPM model. **D-E** Train, testing, and validation cluster penalized regression model. **G-J** Train, testing, and validation full penalized regression model.

**S.Figure2:**
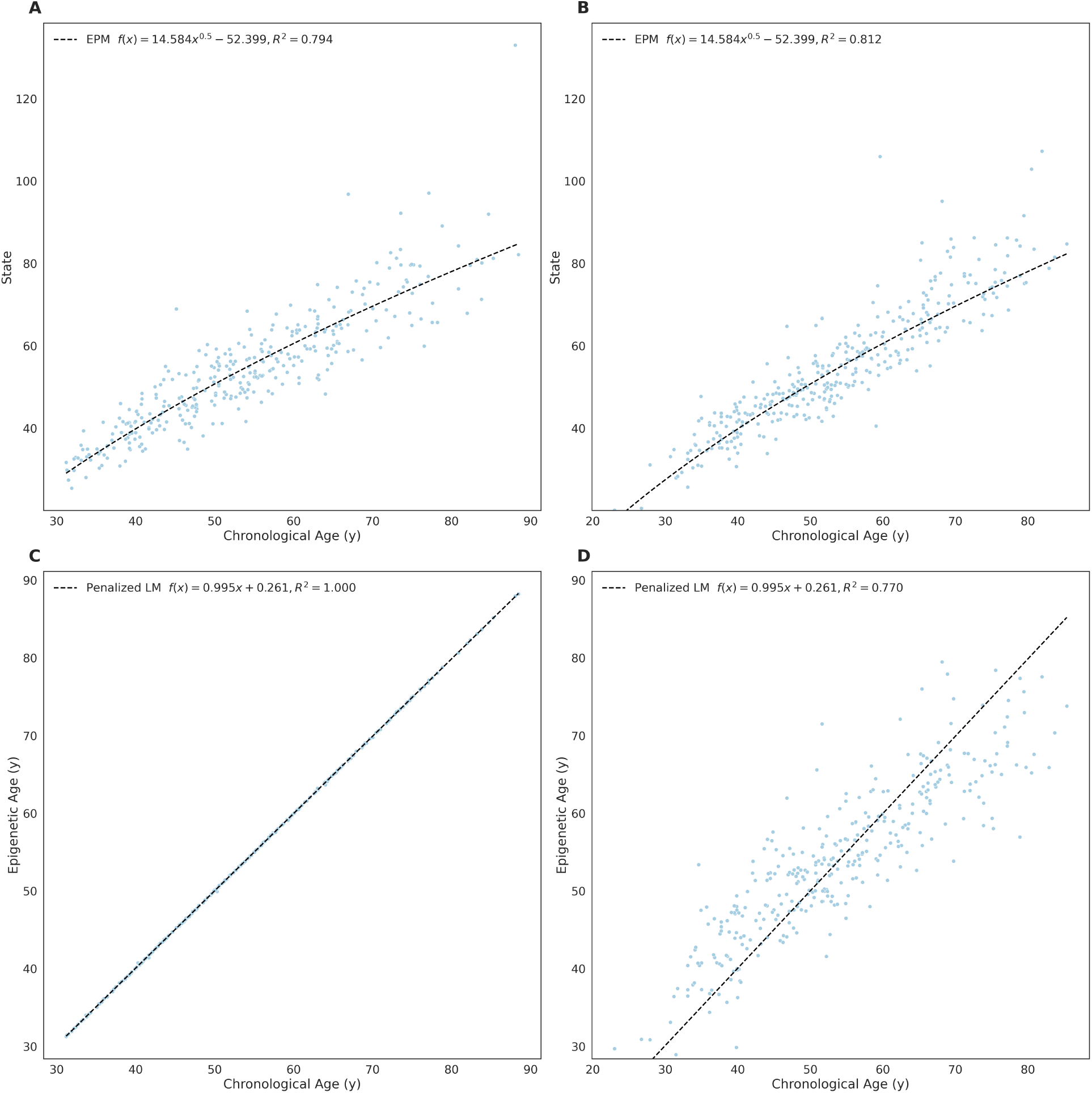
PBB EPM and regression models. **A - B** Train and testing EPM model. **C-D** Train and testing penalized regression model.

## References

1. Horvath, S. & Raj, K. DNA methylation-based biomarkers and the epigenetic clock theory of ageing. en. Nat. Rev. Genet. 19, 371–384 (June 2018).

2. Horvath, S. DNA methylation age of human tissues and cell types 2013.

3. Hannum, G. et al. Genome-wide methylation profiles reveal quantitative views of human aging rates. en. Mol. Cell 49, 359–367 (Jan. 2013).

4. Marioni, R. E. et al. DNA methylation age of blood predicts all-cause mortality in later life. en. Genome Biol. 16, 25 (Jan. 2015).

5. Perna, L. et al. Epigenetic age acceleration predicts cancer, cardiovascular, and all-cause mortality in a German case cohort 2016.

6. Dugué, P.-A. et al. DNA methylation-based biological aging and cancer risk and survival: Pooled analysis of seven prospective studies. en. Int. J. Cancer 142, 1611–1619 (Apr. 2018).

7. Huang, R.-C. et al. Epigenetic Age Acceleration in Adolescence Associates With BMI, Inflammation, and Risk Score for Middle Age Cardiovascular Disease. en. J. Clin. Endocrinol. Metab. 104, 3012–3024 (July 2019).

8. Armstrong, N. J. et al. Aging, exceptional longevity and comparisons of the Hannum and Horvath epigenetic clocks. en. Epigenomics 9, 689–700 (May 2017).

9. Horvath, S. et al. Decreased epigenetic age of PBMCs from Italian semi-supercentenarians and their offspring. en. Aging 7, 1159–1170 (Dec. 2015).

10. Horvath, S. et al. Obesity accelerates epigenetic aging of human liver. en. Proc. Natl. Acad. Sci. U. S. A. 111, 15538–15543 (Oct. 2014).

11. Zhang, Q. et al. Improved precision of epigenetic clock estimates across tissues and its implication for biological ageing 2019.

12. Snir, S., Farrell, C. & Pellegrini, M. Human epigenetic ageing is logarithmic with time across the entire lifespan. en. Epigenetics 14, 912–926 (Sept. 2019).

13. Snir, S., vonHoldt, B. M. & Pellegrini, M. A Statistical Framework to Identify Deviation from Time Linearity in Epigenetic Aging. en. PLoS Comput. Biol. 12, e1005183 (Nov. 2016).

14. Farrell, C., Snir, S. & Pellegrini, M. The Epigenetic Pacemaker: modeling epigenetic states under an evolutionary framework. en. Bioinformatics 36, 4662–4663 (Nov. 2020).

15. Marabita, F. et al. Author Correction: Smoking induces DNA methylation changes in Multiple Sclerosis patients with exposure-response relationship. en. Sci. Rep. 8, 4340 (Mar. 2018).

16. Ventham, N. T. et al. Integrative epigenome-wide analysis demonstrates that DNA methylation may mediate genetic risk in inflammatory bowel disease. en. Nat. Commun. 7, 13507 (Nov. 2016).

17. Tan, Q. et al. Epigenetic signature of birth weight discordance in adult twins. en. BMC Genomics 15, 1062 (Dec. 2014).

18. Johnson, R. K. et al. Longitudinal DNA methylation differences precede type 1 diabetes. en. Sci. Rep. 10, 3721 (Feb. 2020).

19. Voisin, S. et al. Many obesity-associated SNPs strongly associate with DNA methylation changes at proximal promoters and enhancers 2015.

20. Soriano-Tárraga, C. et al. Epigenome-wide association study identifies TXNIP gene associated with type 2 diabetes mellitus and sustained hyperglycemia. en. Hum. Mol. Genet. 25, 609–619 (Feb. 2016).

21. Dabin, L. et al. Altered DNA methylation profiles in blood from patients with sporadic Creutzfeldt-Jakob disease

22. Horvath, S. & Ritz, B. R. Increased epigenetic age and granulocyte counts in the blood of Parkinson’s disease patients. en. Aging 7, 1130–1142 (Dec. 2015).

23. Kurushima, Y. et al. Epigenetic findings in periodontitis in UK twins: a cross-sectional study 2019.

24. Zannas, A. S. et al. Epigenetic upregulation of FKBP5 by aging and stress contributes to NF-κB–driven inflammation and cardiovascular risk. en. Proc. Natl. Acad. Sci. U. S. A. 116, 11370–11379 (June 2019).

25. Braun, P. R. et al. Genome-wide DNA methylation comparison between live human brain and peripheral tissues within individuals. en. Transl. Psychiatry 9, 47 (Jan. 2019).

26. Demetriou, C. A. et al. Methylome analysis and epigenetic changes associated with menarcheal age. en. PLoS One 8, e79391 (Nov. 2013).

27. Tserel, L. et al. Age-related profiling of DNA methylation in CD8+ T cells reveals changes in immune response and transcriptional regulator genes. en. Sci. Rep. 5, 13107 (Aug. 2015).

28. Barrett, T. et al. NCBI GEO: archive for functional genomics data sets— update. en. Nucleic Acids Res. 41, D991–D995 (Nov. 2012).

29. Houseman, E. A. et al. DNA methylation arrays as surrogate measures of cell mixture distribution. en. BMC Bioinformatics 13, 86 (May 2012).

30. Aryee, M. J. et al. Minfi: a flexible and comprehensive Bioconductor package for the analysis of Infinium DNA methylation microarrays. en. Bioinformatics 30, 1363–1369 (May 2014).

31. Johansson, A., Enroth, S. & Gyllensten, U. Continuous Aging of the Human DNA Methylome Throughout the Human Lifespan. en. PLoS One 8, e67378 (June 2013).

32. Liu, Y. et al. Epigenome-wide association data implicate DNA methylation as an intermediary of genetic risk in rheumatoid arthritis 2013.

33. Butcher, D. T. et al. CHARGE and Kabuki Syndromes: Gene-Specific DNA Methylation Signatures Identify Epigenetic Mechanisms Linking These Clinically Overlapping Conditions. en. Am. J. Hum. Genet. 100, 773–788 (May 2017).

34. Dámaso, E. et al. Comprehensive Constitutional Genetic and Epigenetic Characterization of Lynch-Like Individuals. en. Cancers 12 (July 2020).

35. Frey, B. J. & Dueck, D. Clustering by passing messages between data points. en. Science 315, 972–976 (Feb. 2007).

36. Fries, G. F. The PBB episode in Michigan: an overall appraisal. en. Crit. Rev. Toxicol. 16, 105–156 (1985).

37. Safe, S. Polychlorinated biphenyls (PCBs) and polybrominated biphenyls (PBBs): biochemistry, toxicology, and mechanism of action. en. Crit. Rev. Toxicol. 13, 319–395 (1984).

38. Jacobson, M. H. et al. Serum Polybrominated Biphenyls (PBBs) and Polychlorinated Biphenyls (PCBs) and Thyroid Function among Michigan Adults Several Decades after the 1973–1974 PBB Contamination of Livestock Feed 2017.

39. Curtis, S. W. et al. Thyroid hormone levels associate with exposure to polychlorinated biphenyls and polybrominated biphenyls in adults exposed as children. en. Environ. Health 18, 75 (Aug. 2019).

40. Terrell, M. L., Rosenblatt, K. A., Wirth, J., Cameron, L. L. & Marcus, M. Breast cancer among women in Michigan following exposure to brominated flame retardants. en. Occup. Environ. Med. 73, 564–567 (Aug. 2016).

41. Hoque, A. et al. Cancer among a Michigan cohort exposed to polybrominated biphenyls in 1973. en. Epidemiology 9, 373–378 (July 1998).

42. Curtis, S. W. et al. Exposure to polybrominated biphenyl (PBB) associates with genome-wide DNA methylation differences in peripheral blood. en. Epigenetics 14, 52–66 (Jan. 2019).

43. Pidsley, R. et al. Critical evaluation of the Illumina MethylationEPIC BeadChip microarray for whole-genome DNA methylation profiling. en. Genome Biol. 17, 208 (Oct. 2016).

44. Thompson, M. J. et al. A multi-tissue full lifespan epigenetic clock for mice. en. Aging 10, 2832–2854 (Oct. 2018).

45. Thompson, M. J., vonHoldt, B., Horvath, S. & Pellegrini, M. An epigenetic aging clock for dogs and wolves. en. Aging 9, 1055–1068 (Mar. 2017).

46. Pinho, G. M. et al. Hibernation slows epigenetic aging in yellow-bellied marmots en. Mar. 2021.

47. Larison, B. et al. Epigenetic models predict age and aging in plains zebras and other equids en. Mar. 2021.

48. Harris, C. R. et al. Array programming with NumPy. en. Nature 585, 357–362 (Sept. 2020).

49. Pedregosa, F. et al. Scikit-learn: Machine Learning in Python. J. Mach. Learn. Res. 12, 2825–2830 (2011).

50. Seabold, S. & Perktold, J. Statsmodels: Econometric and statistical modeling with python in Proceedings of the 9th Python in Science Conference 57 (2010), 61.

51. Triche Jr, T. J., Weisenberger, D. J., Van Den Berg, D., Laird, P. W. & Siegmund, K. D. Low-level processing of Illumina Infinium DNA Methylation BeadArrays. en. Nucleic Acids Res. 41, e90 (Apr. 2013).

52. Basu, A. Reproducible research with jupyter notebooks

53. Varoquaux, G. & Grisel, O. Joblib: running python function as pipeline jobs. packages. python. org/joblib (2009).

54. Virtanen, P. et al. SciPy 1.0: fundamental algorithms for scientific computing in Python. Nat. Methods (Feb. 2020).

55. Hunter, J. D. Matplotlib: A 2D Graphics Environment 2007.

56. Waskom, M. seaborn: statistical data visualization. J. Open Source Softw. 6, 3021 (Apr. 2021).

57. McKinney, W. Python for Data Analysis: Data Wrangling with Pandas, NumPy, and IPython en (“O’Reilly Media, Inc.”, Oct. 2012).

58. Da Costa-Luis, C. O. tqdm: A Fast, Extensible Progress Meter for Python and CLI. JOSS 4, 1277 (May 2019).

